# Precopulatory behavior and sexual conflict in the desert locust

**DOI:** 10.1101/190793

**Authors:** Y. Golov, A. Harari, J. Rillich, A. Ayali

## Abstract

Studies of mating and reproductive behavior have contributed much to our understanding of various animals’ ecological success. The desert locust, *Schistocerca gregaria*, is an important agricultural pest. However, knowledge of locust courtship and precopulatory behavior is surprisingly limited. Here we provide a comprehensive study of the precopulatory behavior of both sexes of the desert locust in the gregarious phase, with particular emphasis on the conflict between the sexes. Detailed HD-video monitoring of courtship and mating of 20 locust pairs, in a controlled environment, enabled both qualitative and quantitative descriptions of the behavior. A comprehensive list of behavioral elements was used to generate an eight-step ethogram, depicting from first encounter between the sexes to actual copulation. Further analyses included the probability of each element occurring, and a kinematic diagram based on a transitional matrix. Eleven novel behavioral elements are described in this study, and two potential points of conflict between the sexes are identified. Locust sexual interaction was characterized by the dominance of the males during the pre-mounting stage, and an overall stereotypic male courtship behavior. In contrast, females displayed no clear courtship-related behavior and an overall less organized behavioral sequence. Central elements in the sexual behavior of the females were low-amplitude hind-leg vibration, as well as rejecting males by jumping and kicking. Intricate reciprocal interactions between the sexes were evident mostly at the mounting stage. The reported findings contribute important insights to our knowledge of locust mating and reproductive behavior, and may assist in confronting this devastating agricultural pest.

## Introduction

The desert locust, *Schistocerca gregaria*, is one of the most serious agricultural pests. During outbreaks, swarms may consist in millions of individuals (40-80 million locusts per km^2^; e.g. Singh & Singh 1977; Steedman 1988; EI-Bashir et al. 1993), and the damage to crops can be enormous, as the locusts are able to consume hundreds of tons of vegetation per day (Shaluf 2007). Moreover, according to some estimates, 1/10 of the global human population is affected by this pest (Latchininsky et al. 2011).

Locusts have served as important models in the study of various aspects of insect physiology and behavior (e.g. Burrows 1996; Ayali & Yerushalmi 2010; Ayali & Lange 2010; Ariel & Ayali 2015). However, our knowledge of locust courtship and precopulatory behavior is surprisingly limited. Understanding the mating and reproductive behaviors of a species has a fundamental role in the understanding of its ecological adaptation (Kirkendall 1983; Thornhill & Alcock 1983). Specifically, thorough comprehension of the interactions between the sexes may provide new insights for the development of alternative methods for coping with agricultural pests (Boake et al. 1996; Suckling 2000). This should be achieved by a combination of qualitative descriptions and quantitative analyses-the two complementary components of an ethological study (Kasuya 1983).

A comprehensive study of animal behavior should start with a list of behavioral elements (or ‘units’), followed by their chronological appearance, in order to construct a species-specific **ethogram**. The quantification of behavioral elements needs to be based not only on the appearance of these elements, but also on their frequency, their sequence and the probability of transition. Such an approach can identify the typical elements and key transitions during the behavioral ritual (e.g. Klein & De Araújo 2010). The quantification can be aided by using a Markovian chain, also known as a transition matrix (Castrovillo and Cardé 1980; Haynes and Birch 1984). The knowledge gained may contribute not only to deciphering evolutionary relationships between taxa, as in host-parasite interactions, but also to the understanding of mate recognition and sexual conflicts (Paranjape 1985; Curkovic et al. 2006; Cozzie & Irby 2010; Gaertner et al. 2015), and specifically so in agricultural pests (Walgenbach & Burkholder 1987; Rojas et al. 1990; Wang & Millar 2000; Zahn et al. 2008).

Some aspects of the sexual behavior of *S. gregaria* have been previously addressed (Uvarov 1928; Husain and Mathur 1946; Laub-Drost 1959, 1960 cited in Uvarov 1966, 1977; Popov 1958; Loher 1959, 1961; Pener 1965, 1967a,b; Norris 1964; Odhiambo 1966; Roffey & Popov 1968; Strong & Amerasinghe 1977; Uvarov 1977; Amerasinghe 1978a; Pener & Lazarovici 1979; Inayatullah et al. 1994; Njagi & Torto 2002), but much of the required knowledge is still lacking. The published descriptions and quantifications of the sexual behavior of both sexes are either limited (Strong & Amerasinghe 1977; Inayatullah et al. 1994), too general, or focus predominantly on the male (e.g Pener 1967b; Amerasinghe 1978b). In addition, previous studies suffer from inconsistencies (e.g. different names for similar behavioral element). Finally, little effort has been dedicated to the study of sexual conflict in this insect. The desert locust displays a clear sexual dimorphism in the gregarious phase, with fully mature males being bright yellow and females being beige-brown to yellowish (Chauvin 1941 cited in Pener & Simpson 2009; Norris 1954; Pener 1967b). As is the case for many other Acridids, little is known regarding the means of sexual recognition in the desert locust (Whitman 1990). It is postulated, however, that visual and chemical signals play an important role (Obeng-Ofori et al. 1993, 1994; Franck and Schmidt 1994; Inayatullah et al. 1994; Ely et al. 2006; Seidelmann & Warnstorff 2001). Courtship and mating behaviors can be roughly divided into two sequential stages: pre-copulatory and post-copulatory (with copulation defined as the time when sperm is transferred). The pre-copulatory stage can be divided into two further sub-stages: pre-mounting, comprising all the behavioral elements leading to a successful mounting attempt; and mounting, culminating in successful copulation. Locust courtship is considered simple and primitive (Loher 1959; Uvarov 1966, 1977; Oberlin 1974 cited in Strong & Amerasinghe 1977). As in many grasshoppers, males of *S. gregaria* have been reported to be the dominant gender during the sexual-interactions (Norris 1964; Pener 1965, 1967b; Strong & Amerasinghe 1977; Amerasinghe 1978a; Inayatullah et al. 1994). Briefly, the male’s sexual intention is initially demonstrated through his orientation towards the female, followed by a stealthy slow approach and a surprise attempt to mount her. Once mounting, the male grasps the female using his front and mid-legs. Copulation is achieved when the male moves his abdomen along the side of the female and connection between the genitalia is established. In contrast to the males, gregarious females have been considered to demonstarte no clear courtship behavior (Norris 1964; Pener 1965, 1967b; Strong & Amerasinghe 1977; Amerasinghe 1978a; Inayatullah et al. 1994). Nonetheless, the rejection of courting males has been reported, including the female’s jumping (before and during mounting), kicking, and lateral movements of her abdomen in the attempt to prevent copulation (Loher 1959; Strong & Amerasinghe 1977; Uvarov 1977). Hind leg vibration and wing stridulation have been reported to be displayed during the pre-copulatory behavior (Morse 1896; Norris 1954; Laub-Drost 1959 cited in Uvarov 1977; Loher 1959, 1961; Otte 1970; Uvarov 1966, 1977), as in other acridid grasshoppers (Haskell 1957, 1958; Otte 1977). Unlike wing stridulation (displayed by both sexes), the vibration of the hind legs is soundless and much more common in the female (Loher 1959). The role of both behavioral elements in the sexual interaction has remained uncertain (Loher 1959; Uvarov 1966; Otte 1970).

The major goals of this work were to generate an ethogram, comprising and accompanied by both qualitative and quantitative tools for studying the sexual behavior of the two sexes of the desert locust during the pre-copulatory stage. This included generating a detailed list of all related behavioral elements, and consolidating the relevant terminology (i.e. ‘*nomenclatura*’). The generated ethogram includes all the behavioral elements, their occurrences, and their sequence during the sexual interaction. This had enabled an elaborate description of the conflict between the sexes in gregarious locusts. In an accompanying study (Golov et al., in preparation), we employ the tools developed herein for a comparative investigation of the two density-dependent locust phases.

## Material & Methods

### Animals

Desert locusts, *Schistocerca gregaria*, from our colony at Tel Aviv University (Ayali et al., 2002) were reared for many consecutive generations under crowded conditions (i.e. approaching the gregarious phase), 100-160 individuals in 60 aluminum cages. All cages were located in a dedicated room under a constant temperature (29-31°C) and light cycle of 12: 12 D: L. Supplementary radiant heat was supplied during day-time by incandescent 25 W electric bulbs, resulting in a day temperature of c. 37°C. Locusts were provided daily with fresh wheat and dry oats, and plastic caps (300cc) filled with moist sand for oviposition.

All locust individuals in the experiments were adult virgin males and females. Virgin adults were obtained by marking newly-emerged adults with non-poisonous acrylic paint within 24 hours following ecdysis. Males and females were separated into single sex “cohort cages” every 3 days. Thus, in each cohort cage the maximum age range of the individual locusts was less than 72 hours. The cages were maintained under the same rearing conditions as above. For the observations we used 12-14-days-old males, when their yellowish coloration had reached stage V (see Norris 1954 & Loher 1961). This stage is known to coincide with sexual maturity. Females were 18-20 day-old, sexually mature, based on our preliminary work and other previous reports (Hamilton 1955; Injeyan & Tobe 1981; Mahamat et al. 1993; Wybrandt & Andersen 2001; Ould Ely et al. 2006; Nishide and Tanaka 2012). Only fully intact insects participated in the observations.

### Experimental design

Experiments were carried out in an isolated room, with temperature and light conditions similar to that in the rearing room. A plastic observation cell (14x13x24 cm) was initially divided, by an opaque plastic partition into two compartments, to separately host the male and the female. The sensitivity of *S. gregaria* color vision is mainly in the very short wavelengths of both UV and blue (320 and 450·nm), and to a lesser extent, also in the green range (light 530·nm) (Eggers & Gewecke 1993; Schmeling et al. 2014). Hence, until initiation of the experiment the cell was illuminated by a red light (to reduce the insects’ stress). Five minute after placing each locust (one male and one female) into its own compartment, the experiment was initiated by carefully removing the partition between the compartments, and replacing the red light with two regular 25 W light bulbs. Two identical observation cells, separated only by a dens plastic mesh (not sealed), were used simultaneously (each housing one pair of locusts), generating crowd-like conditions by allowing the flow of auditory, olfactory and visual cues. Experiments lasted 3 hours, or until copulation had occurred, if earlier, and were recorded by a SONY HDR-PJ820E video camera.

Two rounds of experiments were carried out daily: at 08:00 AM and 15:00 PM. Out of an overall 31 monitored experiments, 20 ended in copulation within the defined time of 3 hours, and were used in the analyses.

In order to further verify the significance of the females’ active rejection behaviors (jumping and kicking) and their tentative role in female choice, a separate series of experiments were carried out. Here we examined male mating success when facing “handicapped” females. The rejection attempts of these females were constrained by means of a small rubber band confining the hind legs’ femur and tibia in a folded position, and thus, preventing the female from either jumping or kicking. The number of male mounting attempts and successful mounts were compared between pairs of males and constrained females (N=10) and males and unrestrained females (N=20).

### Data analyses

The recorded videos of the behavior of each pair were reviewed and analyzed using J-watcher software (version 0.9 for Windows).

Behavioral elements were identified in order to construct the locusts’ pre-copulatory behavior. These included both repetitive (lengthy, e.g. the vibration of the hind leg femur) and discrete (momentary, e.g. jumping) behaviors. The two behavioral types were counted, with a ‘count’ relating to the duration of a behavior from initiation until termination. Behavioral measurments were taken only if the male and female were at a distance of less than 10 cm (i.e. an ‘encounter’). For both pre-mounting and mounting behaviors, the following parameters were measured and compared for both sexes: (1) In order to obtain the pattern or chronological sequence of the behavioral repertoire, the relative time to initiation of each behavior was noted (relative to the total time of the relevant stage, either pre-mounting or mounting). (2) The probability of a specific behavior occurring (PO=1 if the behavior occurred at least once, and 0 otherwise). (3) The frequency of occurrences of a specific behavioral element.

A kinematic diagram was constructed, based on a first-order Markov model, for all the transitions between pairs of behavioral elements (i.e. preceding—following elements) that are mutually exclusive (Baker & Carde 1979). All the behavioral elements in this analysis were considered nodes and used to construct a transitional matrix. The transition probability (TP, also known as ‘conditional probabilities’; Wood et al. 1980) was first calculated based on all possible transitions between a pair of nodes in the matrix, for each experiment (see also Brown 1974; Leonard and Ringo 1978; Markow and Hanson 1981). Next, the average of each transition was calculated among all 20 pairs for each sex (following the method described by Charlton & Cardé 1990). Self-transitions were scored as structural zeroes (Baker & Cardé 1979), and impossible transitions were left blank (Haynes & Birch 1984). Those behavioral elements that were not mutually exclusive with any of the other elements (‘antennal movement’, ‘palp vibration’, ‘genital-opening’, ‘abdominal wagging’) were excluded from this analysis. Thus, the behavioral transitional matrix comprised 25 elements for the male, and 18 for the female. Transitional probabilities (i.e. TP) ≤10% are not presented.

Most of the statistical output and data analysis were conducted in GraphPad Prism version 6.04 for Windows, JMP®, Version 12.0.1 SAS Institute, and some in Matlab (MathWorks, USA Inc.) and Canvas draw 2.0 (Deneba Systems, Miami, FL).

## Results

### The sexual behavior of the desert locust

As noted above, the behavioral elements that lead to copulation (i.e. those that can be identified during the pre-copulatory phase), can be divided into two stages: Table 1 lists all the elements comprising the pre-mounting stage, and Table 2 lists all the elements comprising the mounting stage (ending in copulation). Among the behavioral repertoire listed in Table 1 and 2, several elements have been described previously. However, those descriptions tend to be episodic, with different authors providing different descriptions for the same behavior, or referring to the same behavior by different names, etc. Eleven elements are novel, and are described here for the first time.

**Table 1.** The behavioral repertoire during the pre-mounting stage.

| Behav. Element | Description | Sex |  | Citation |
| --- | --- | --- | --- | --- |
|  |  | m | f |  |
| Grooming | Eyes, antennas, mouth part, mid and front legs, wings and abdominal grooming | + | + | O'Shea 1970; Rowell 1971; Uvarov 1977; Berkowitz & Laurent 1996 |
| Antennal movement | Movement of the antennas | + | + | Loher 1959; Wallace 1959 |
| Palp vibration | Vibration movements of both labial and maxillary palps | + | + | Uvarov 1977 |
| Searching | Combination of walking, scanning, antennal movement and palps vibration | + | + | based on Inayatullah et al. 1996 |
| Orienting | Anterior side is directed towards the female | + | - | Otte 1970; Amerasinghe 1978 |
| High hind leg (HL) femoral vibration | Femur is lifted to a perpendicularly position relatively to the ground (~90°) while vibrating the hind legs. | + | + | Loher 1959 |
| Low hind leg (HL) femoral vibration | Femur is lifted to a perpendicularly position relatively to the ground (<90°) while vibrating the hind legs. | + | + | Loher 1959 |
| Wing flapping | Flapping movements of both wing pairs | + | + | Wies-Fogh 1956a,b cited in Uvarov 1966; Uvarov 1977 |
| Long stridulation | Rapid vibration of the wing-pairs producing whizzing noise | + | - | Loher 1959 |
| Short stridulation | Wings beating against | + | - | Loher 1959 |
| Abdomen wagging | Wagging (mostly lateral) movements of the abdomen | + | + | * |
| Genital opening | Rhythmic opening of the genital-opening | + | + | * |
| Initiating physical contact | Locusts touching each other | + | + | Popov 1958 |
| ***Mutual antennal contact*** | Locust antennas touching each other | + | + | * |
| Slow repeating hind leg elevation | Slow elevation of the hind legs | + | - | * |
| Approaching | Walking clearly directed towards other sex | + | + | Popov 1958; Loher 1959 |
| Walking away from the male | Distancing from the male by walking | - | + | Popov 1958; Loher 1959 |
| Jumping away from the male | Distancing from the male by jumping | - | + | Popov 1958; Loher 1959 |
| Mounting by Climbing | Attempting to mount the female by climbing | + | - | * |
| Mounting attempt by Jumping | Attempting to mount the female by jumping | + | - | Uvarov 1928; Husain & Mathur 1946 |

Behaviors which are shared and mutually exhibited by both sexes are presented in italic bold font. \* Represents elements which are described for the first time.

**Table 2.** The behavioral repertoire during the mounting stage.

| Behav. Element | Description | Sex |  | Citation |
| --- | --- | --- | --- | --- |
|  |  | m | f |  |
| ***Mounting*** | The male mounting the female/ female is being mounted by the male | + | + | Uvarov 1928; Husain & Mathur 1946 |
| Avoidance | Elevation of the two hind legs femur close together in the air | + | - | * |
| Blocking | Lifting both Hind legs in the air perpendicularly to the ground, femur and tibia folded | + | - | * |
| Jumping | Jumping while carrying the male | - | + | Loher 1959 |
| Kicking the male | Directed kicking towards the male | - | + | Loher 1959 |
| Abdominal bending | Abdominal tips and genital are bended sideways | - | + | Strong & Amerasinghe 1977 |
| Abdomen grounding | Abdomen is pressed to the ground | - | + | * |
| ***Male's dislodgement*** | Male is dislodged from the female's back | + | + | Popov 1958; Uvarov 1977 |
| Lifting attempt | Males attempt to lift the female by pushing against the ground and straightening of the hind legs | + | - | * |
| Antennal movement | Movement of the antennae | + | + | Uvarov 1977 |
| Palp vibration | Vibration of both labial and maxillary palps | + | + | Loher 1959 |
| Hind legs vibration | Femur is being positioned perpendicularly to the ground while tibia is flexed 90° to the femur. | + | - | Loher 1959 |
| Short Stridulation | Wings beating against each other resulting in short and sharp sounds | + | - | Loher 1959 |
| Long stridulation | Rapid vibration of the wings while in resting position | + | - | Loher 1959 |
| Lateral hind leg vibration | Lateral vibration of the hind legs femur | - | + | Loher 1959 |
| Hind leg grounding | Pushing the hind legs against the ground. | + | - | * |
| Genital opening | Rhythmic opening-closing of the genital opening | + | + | * |
| Copulation attempt | Male's abdomen is placed to the side of the female's ones in order to reach her genitalia | + | - | Loher 1959 |
| ***Copulation*** | Copulation defined as the time when sperm is transferred |  |  | Pener 1967 |

Behaviors which are shared and mutually exhibited by both sexes are presented in italic bold font. \* Represents elements which are described for the first time.

The probability of each element being demonstrated varies greatly. Figure 1 denote the probability of a behavioral element occuring (PO), separately for males and females for the pre-mounting and mounting stages. The behavioral elements appear in a consecutive order and are grouped following a further subdivision: S1-S7, from initiation (S1) to copulation attempt (S7), culminating in S8, copulation. Figure 1 presents the elements that involve all body parts (denoted by different colors), including legs, wings, palps and antenna, and abdomen. Some of the sub-stages are characterized by a consistently high PO (e.g. S1 during the pre-mounting; Fig. 1) while that of others varies greatly. Moreover, the PO of the elements demonstrated by the male or the female within the same sub-stage differs (e.g. compare S1-2 or S5-7 in Fig. 1). Generally speaking, a high PO reflects the importance of a behavioral element within the overall sequence. However, there may be low PO elements that nonetheless have a crucial functional significance: e.g. those instrumental in inter‐ and probably also intra-sexual communication (e.g. leg vibration, wing flutter and stridulation). Illustrations of the different behavioral elements are provided in figure 2.

**Figure 1.**
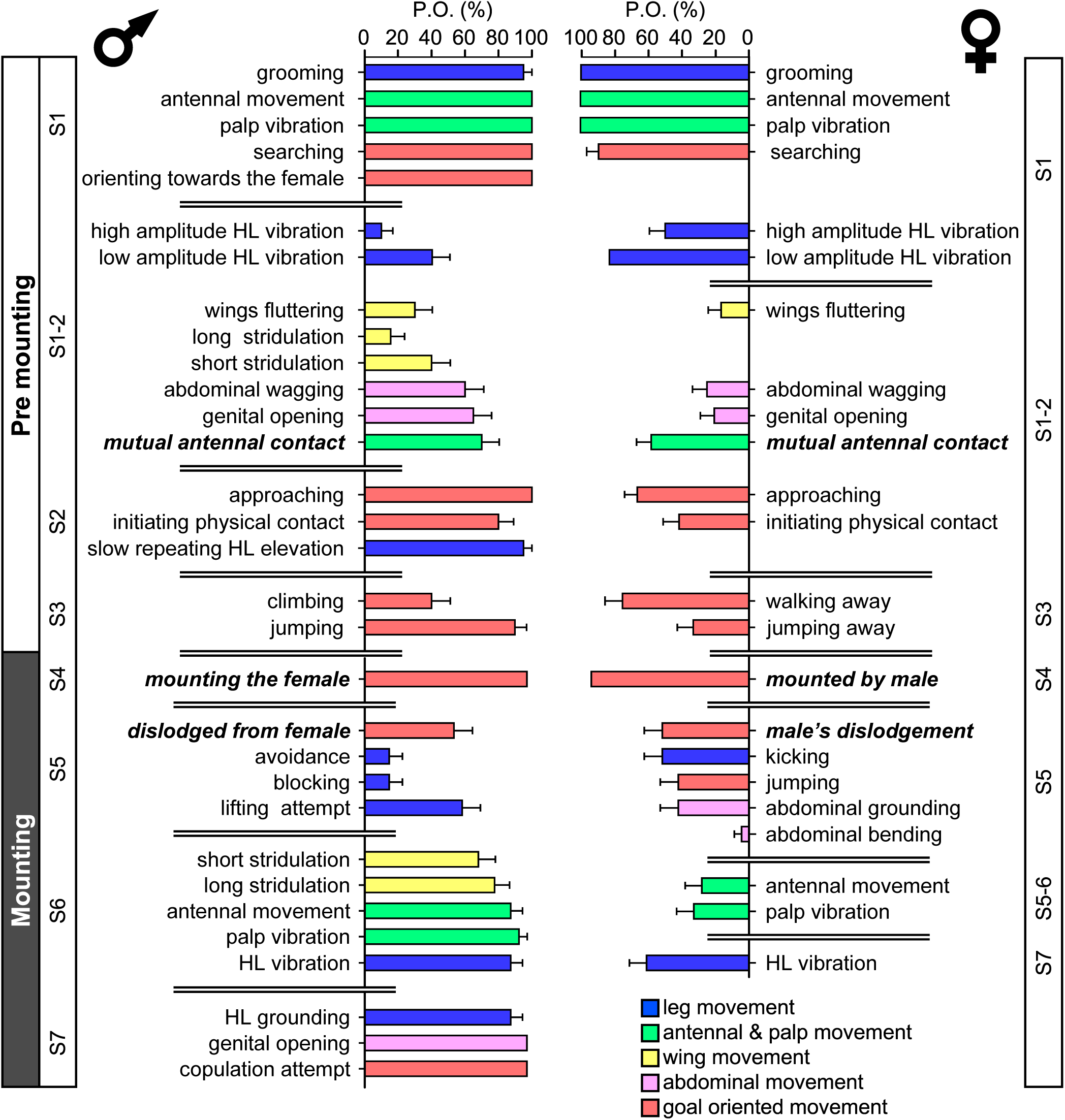
The precopulatory behavioral repertoire of the male (left) and female (right) desert locust. The pre-mounting and mounting behavioral elements are listed from step 1 to 7 (S1-S7) and color coded according to relevant body part. The probability of an element occuring (PO %; mean + SEM) is shown. Behavioral elements that are shared and mutually exhibited by both sexes are presented in italic bold font.

**Figure 2.**
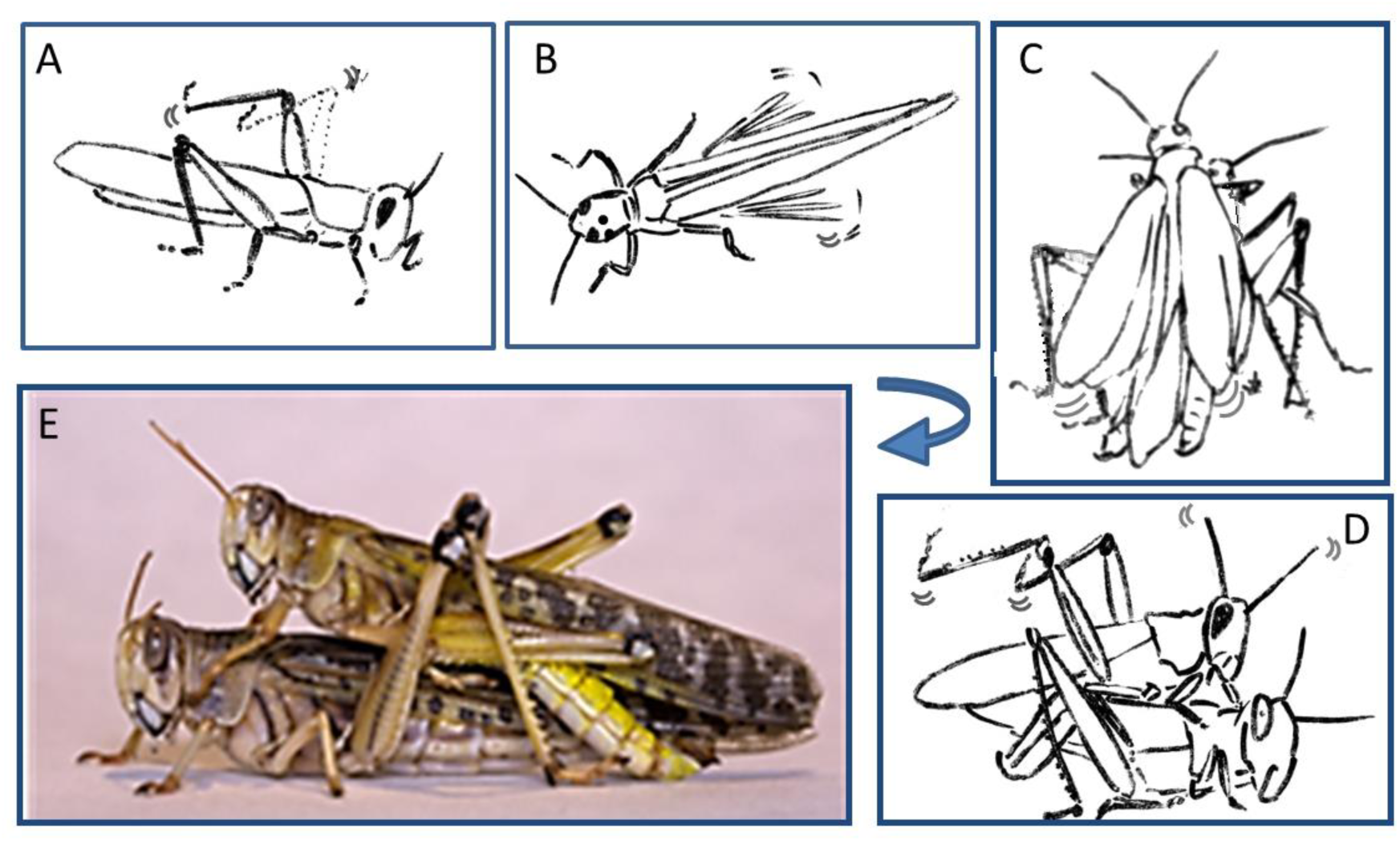
Representative behavioral elements observed during the pre-mounting (A-B) and mounting (C-E) stages: A. Male slow repeated hind leg elevation. B. Female hind leg low and high-amplitude vibration. C. Male short and long wing stridulation. D. Male hind leg vibration and copulation attempt. E. Successful copulation. The animations in A-D were drawn from images taken from video sequences.

An ethogram was constructed (Fig. 3) in order to better characterize the behavioral sequence comprising the pre-copulatory behavior. The ethogram provides the pre-mounting and mounting stages (consistent with Fig. 1), presenting them as an ordered, hierarchical flow-chart. This representation also allowed us to include and emphasize junctions or decision points (denoted by the traffic lights in Fig. 3). These junctions represent the culmination of the conflict between the sexes, e.g. a point at which the female was successful in preventing a mounting attempt by jumping away, or a point at which the male was thrown off the female’s back. Illustrations of behavioral elements of an antagonistic nature can be seen in Figure 4.

**Figure 3.**
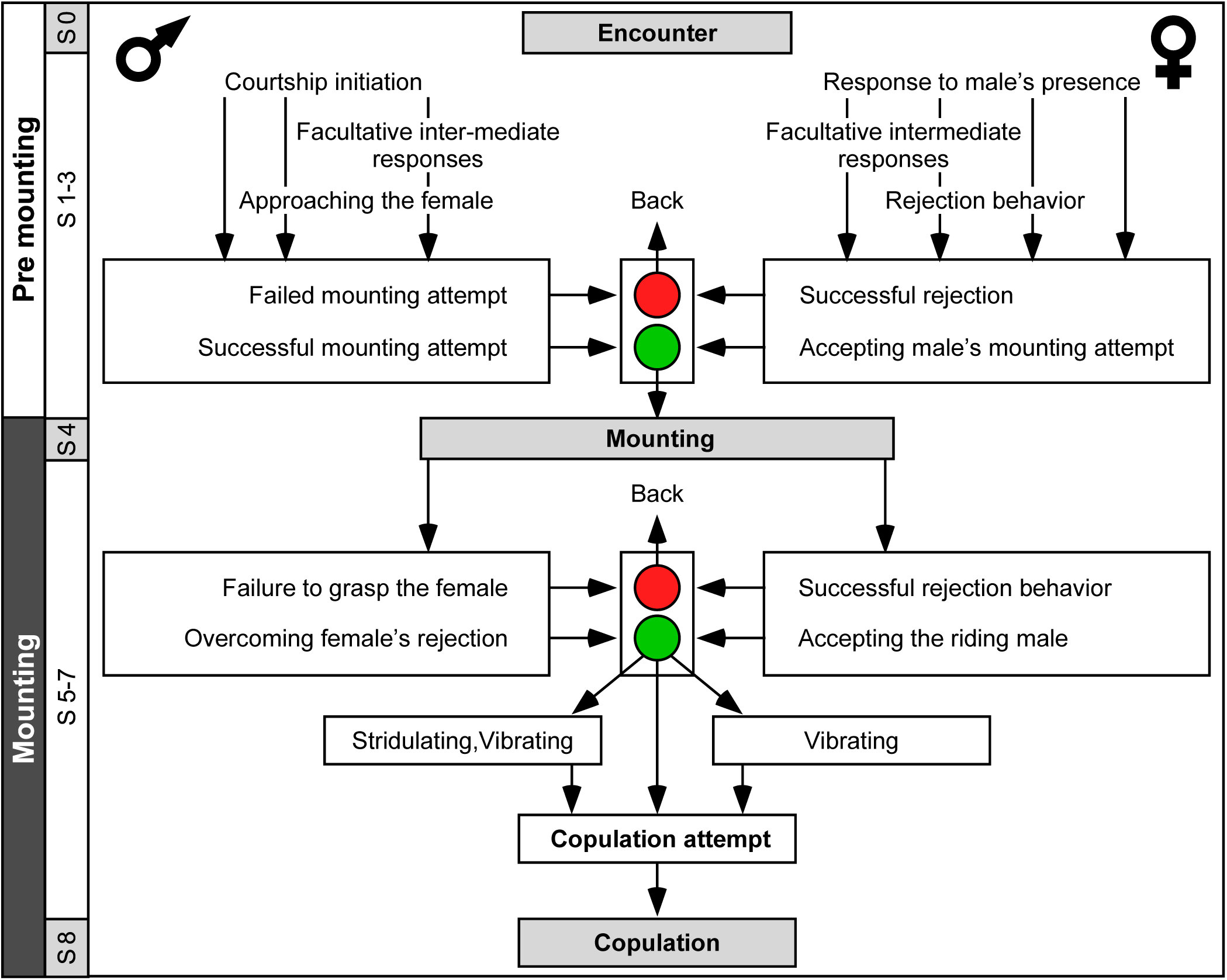
An ethogram depicting the desert locusts pre-copulatory interactions leading to copulation. The male behavior is on the left, and that of the female on the right. S1-S8 indicate the chronological step number during the pre-mounting and mounting stages. Traffic lights denote points at which female choice takes place (steps 3 and 6); Red is associated with rejection of the male. Green is associated with the female tolerating the male.

**Figure 4.**
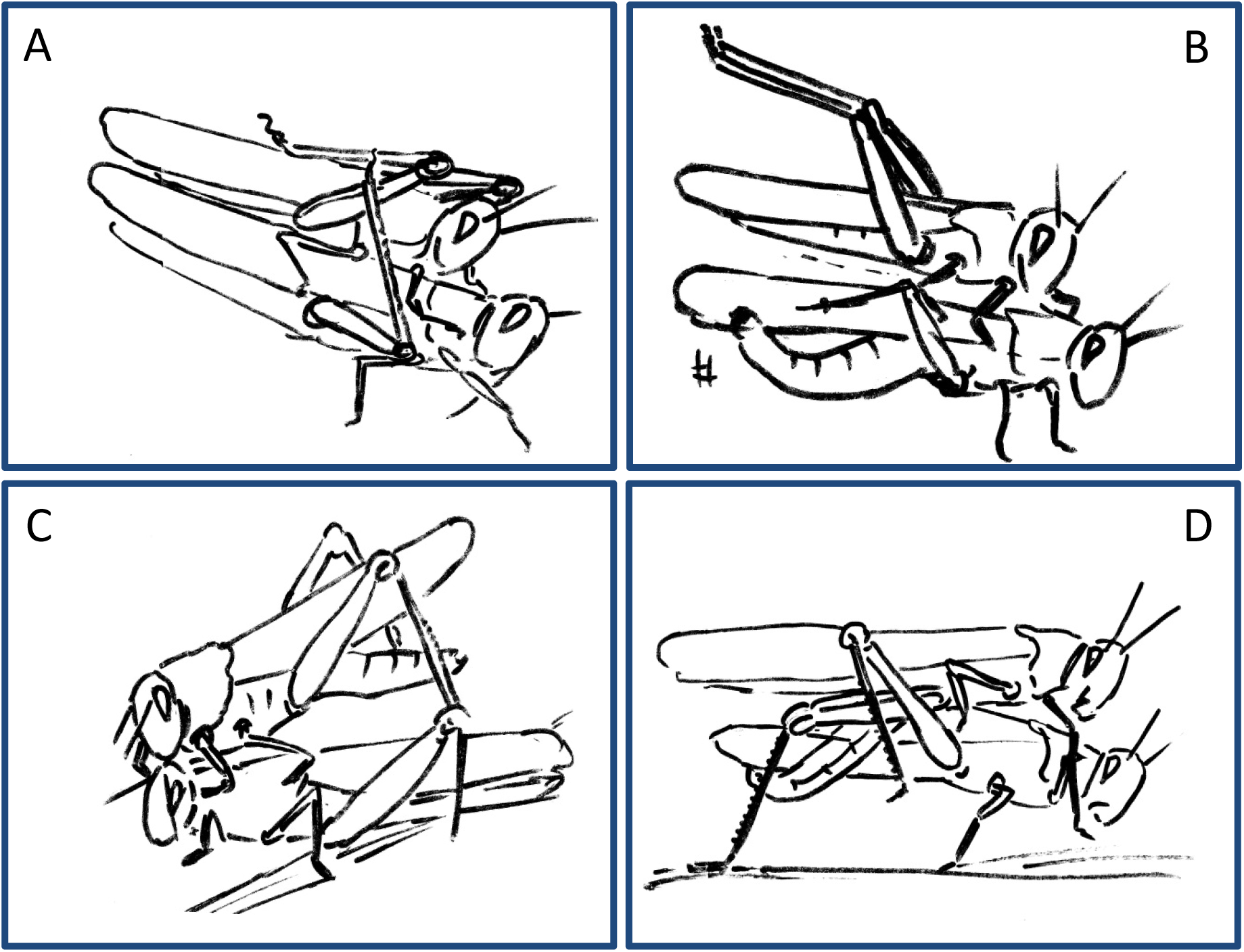
Examples of female rejection behaviors and male responses during the second point of mate choice (second traffic light in the ethogram in Fig. 3). A. Male attempts to block the female’s kicking using his hind legs. B. Female displaying lateral abdomen bending behavior while also kicking, and male responding to kicking by avoidance behavior. C. Female pressing her abdomen to the ground to avoid mating (i.e. ‘abdominal grounding’). D. Male managing to mate with the female by pushing with his hind legs and lifting her.

Further information regarding the flow of the behavioral elements and the overall sequence of the behavior can be obtained by also including, beyond the ordered description of the elements, the probability of a transition from one element to the other. This approach regards the behavioral sequence as a Markov process or Markov chain, in which the appearance of each behavioral element affects or predicts the probability of the appearance of another. Figures 5 and 6 use a similar color code as that presented in Figure 1 to indicate the different behavioral elements constituting the sub-stages (S1-8), presented in Figures 1 and 3. These kinematic diagrams denote a weighted directed network composed of the above introduced different behavioral elements presented by males (Fig. 5) and females (Fig. 6), where the weights are the transition probabilities (TP). As can be seen, this method of presentation clearly discriminates between behavioral elements constituting the relatively consistent or major trunk (depicted 0-8 in Fig. 5, and 0-5 in Fig. 6), as well as the various possible detours or diversifications from it. It also serves to highlight several sex-specific characteristics, as discussed below.

**Figure 5.**
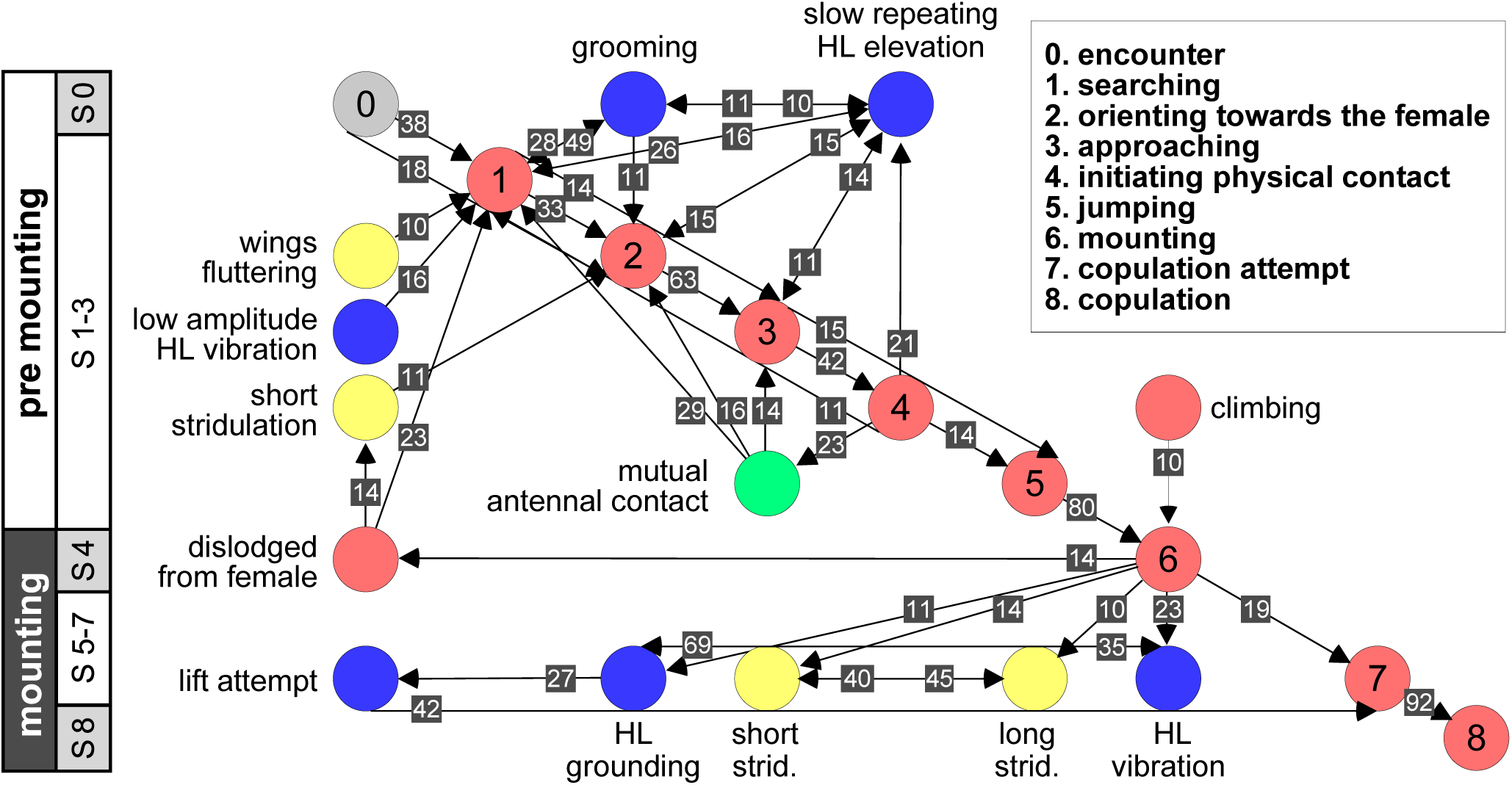
A kinematic diagram depicting the pre-copulatory behavior of male locusts (N=20); arrows represent transitions between behavioral elements. The numbers on a gray background denote the mean transitional probability (TP) between each pair of behavioral elements. Two way transitions are depicted by double-headed arrows (numbers relate to the closer arrow head). The color of the circles representing the different behavioral elements corresponds to the color index used in Figure 1. The different steps in the pre-mounting and mounting stages are noted.

**Figure 6.**
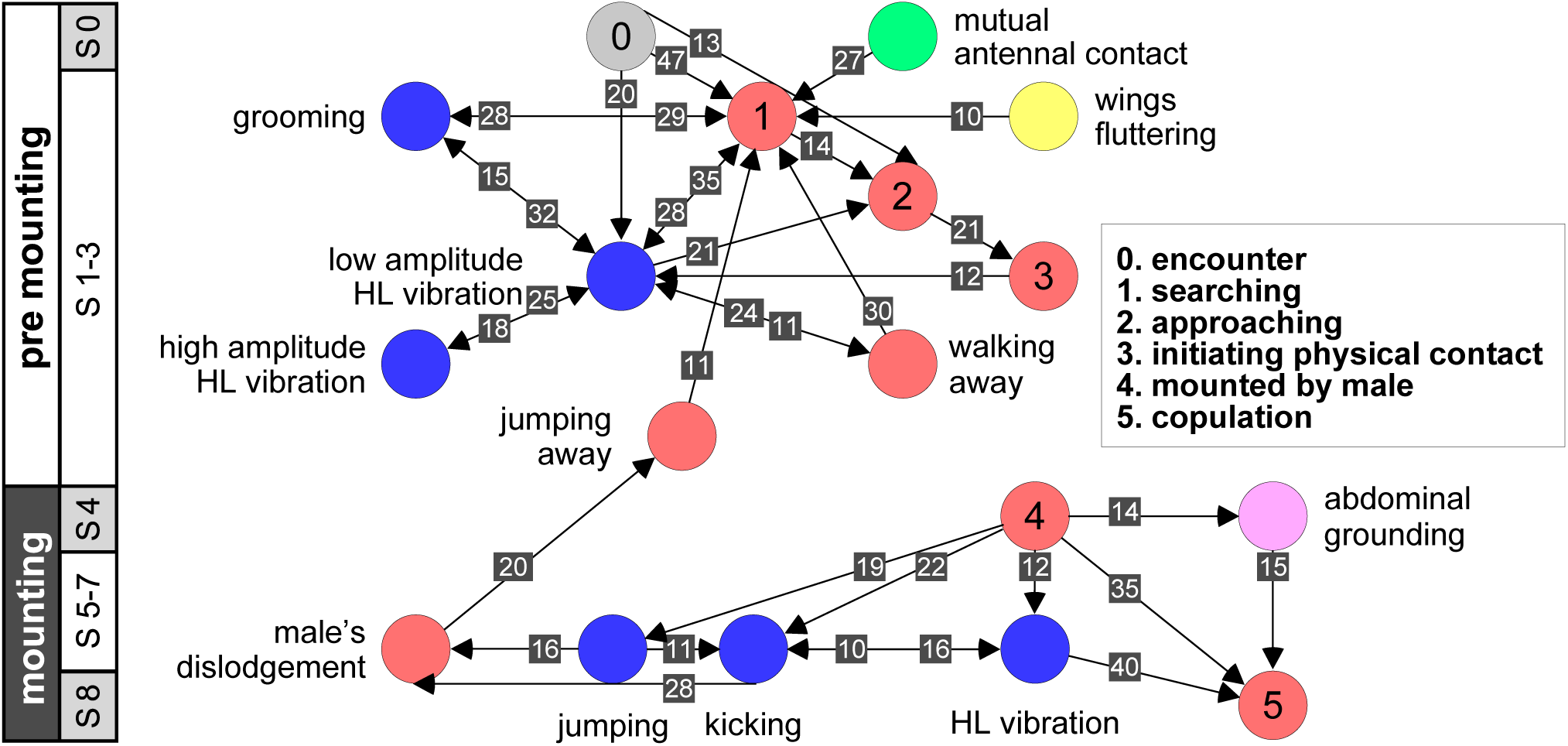
A kinematic diagram depicting the sexual behavior of female locusts (N=20); details as in figure 5.

In the following we provide further details of certain male‐ and female-specific behavioral elements, as well as further insights into the conflict between the sexes.

### Sex-specific sexual behaviors and conflict between the sexes

The strategy employed by males during pre-mounting can be described as stalking, pursuit and attack. Overall courtship in our experiments was somewhat limited. Upon identifying the female, the male commonly demonstrated ‘high-stepping walking’ behavior, carrying his body high above the ground. In some cases, this was intensified prior to jumping in an attempt to mount the female, to an extent that his front legs were raised in the air. Increased self-grooming was shown by all males (PO=100%); males groomed the antennae, the compound eyes, the front or mid pairs of legs and the posterior part of the abdomen. Several behavioral elements, commonly shown during pre-mounting, are described here for the first time. These comprise: lateral wagging movements of the abdomen (‘abdominal wagging’), repeated extension movements of the subgenital-plate and the epiproct (‘genital-opening’), and repeated slow elevation of the hind legs. The latter was performed by most males (PO=95±5%) just after (TP=14%) or before (TP=11%) approaching the female.

Once successful in mounting the female’s back, mostly via jumping, the majority of males were quick to cling to the lateral sides of her pronotum socket (or its edges; see Fig. 2D and E) in order to adjust their grip. Stridulation and hind leg vibrations were more frequent during mounting than pre-mounting, although the cumulative time of mounting (1.65 ± 0.41 min) was much shorter than in pre-mounting (64.23 ± 10.93 min).

The females’ overall sequence of behavioral elements was much less stereotypic compared to that of the males (as also evident from Figs 1, 5 and 6). In spite of the dominant part played by males, the first indication of encounter was usually demonstrated by females (17 out of 20 pairs). Hind leg vibration was a characteristic element of females pre-copulatory behavior, as demonstrated by the high values of both PO and TP (Fig 1 and 6). However the most dominant behavioral feature was the female rejection of the males (Fig. 4).

During the pre-mounting stage, female rejection was displayed by either jumping or walking away from the male. ‘Walking away’ (PO 75%) was commonly followed by the lower amplitude hind leg vibration (TP=24%). The most common rejection element during the mounting stage was kicking (PO=55%).

Both kicking and jumping often caused the mounted male to lose his grip and dislodge from the females back. In fact, more than half of the mated males were dislodged from the female (PO=55%). Females also exhibited “passive rejection” elements, including pressing the abdomen against the ground and thus preventing the male from inserting his abdomen below hers (‘abdominal grounding’ described here for the first time; Fig 4C) and less frequently lateral abdominal bending (Fig 4B; PO=5%), whichwas very efficient in preventing copulation.

Male behavioral elements that were intended to avoid or overcome female rejection are also described here for the first time. These comprised: attempting to block the female’s kicks with the male’s own hind legs (Fig 4A), and elevation of the hind legs with the tibia extended, while keeping the legs close together, in order to minimize the area exposed to the female’s kicking (avoidance; Fig 4B). Naturally, males occupied with these defensive behaviors could not progress toward copulation. An intriguing newly described element is that of the male’s attempt to overcome the female abdominal grounding behavior by pushing with his hind legs and lifting her up (Fig 4D). This reciprocal interaction is shown in a specific path of transitions in Figure 5-‘mounting’ →-‘grounding of the hind legs’ →-‘lifting attempt’ →‘copulation attempt’.

In order to further explore the selected “female choice” stages (S3 and 5; Fig. 3) and verify the significance of female active rejection behaviors and their effect on the males’ mating success, we prevented females from jumping and kicking by means of a small rubber band over their folded hind legs. This manipulation indeed resulted in no active rejection by the constrained females. Consequently, the number of male mounting attempts on these females was significantly lower than that in the control group (med=1<2, U= 40.50, N_1_=10, N_2_=20; p<0.01), and 100% of the pairs comprising a constrained female and a normal male ended in copulation. The males that mounted constrained females displayed none of the documented defense behaviors (see above).

## Discussion

The sexual behavior of the desert locust *S. gregaria* has been previously addressed in various studies (e.g. Uvarov 1928, 1977; Norris 1964; Pener 1965; Odhiambo 1966; Roffey & Popov 1968; Amerasinghe 1978a; Pener & Lazarovici 1979; Njagi & Torto 2002), and was mostly described as primitive and reduced (Popov 1958; Loher 1959; Strong & Amerasighe 1978b; Pener &f Shalom 1987; Inatullah et al. 1994). None of those studies, however, were dedicated to a synchronized, comprehensive investigation of the behavior of the two sexes and their sexual interactions. Loher (1959) for example, although devoting much effort to describing locust sexual behavior, did not include any quantitative measures of the different behavioral elements. In a first attempt to quantify the pre-copulatory behavior of the male, Pener (1967a,b) employed a measure of “average percentage of time spent on sexual behavior”, but with sexual behavior comprising only copulation, sexual attack, or mounting another locust. In a later report, recording the time spent in sexual behavior, Wajc and Pener (1969) noted the great need for elaborate quantitative methods in the study of the sexual behavior of *S. gregaria*. While other quantification efforts (e.g. Inatullah et al. 1994) presented some accounts of behavioral elements, they provided only limited descriptions of the pre-copulatory behavior in a rather anecdotal manner, and similar to previous work focused mostly on males.

In the current study we provide in-depth data on the pre-copulatory behavior of the desert locust in the gregarious phase, comprising both qualitative descriptions and quantitative measures. A detailed list of behavioral elements is prrsented, incorporating eleven elements that are described here for the first time. An ethogram of the sexual behavior of both sexes, from first encounter until copulation, has enabled us to describe the dynamics of the behavior, including the probability of each element being demonstrated and the transitions between elements. Overall eight distinct steps were identified as comprising the two pre-copulatory stages: pre-mounting (S1-3) and mounting (S4-8). Most importantly, two points of conflict between the sexes were recognized and investigated in depth.

### Male sexual behavior

A major characteristic of locust sexual behavior is that of the males’ dominant role in the courtship ritual (Norris 1964; Strong & Amerasinghe 1977; Inayatullah et al. 1994). Our findings well demonstrate this point: that the overall initiative is always by the male. Upon encountering a female, the males displayed a combination of self-grooming, palp vibration and antennal movements (see also Loher 1959). The latter is a known characteristic of male sexual behavior in the family Acrididae (Pickford & Gillott 1972; Otte 1970; Riede 1987). Onset of the rather limited courtship behavior can be recognized initially by the display of “orienting”, in which the male points his antennae towards the female. This behavior is common in the subfamily Catantopinae (Otte 1970).

Another important feature of the male pre-copulatory behavior is its relative consistency, as suggested by Loher (1959) for the courtship behavior of male grasshopper in all Catantopinae species. This stereotypical nature is evident in the present work from the high values of both the PO and TP quantitative measurments. Orientation was followed by a slow, stealthy approach and a sudden jump in the male’s attempt to mount the female. Upon mounting, the male then displayed various stridulation and vibration behavioral elements, culminating in copulation attempts and copulation.

Overall, male sexual behavior varied more during the pre-mounting than during the mounting sub-stages. This was expressed in both, the larger repertoire of elements and the higher variability of their occurrence (PO). A major behavioral element during pre-mounting was that of the slow elevation of the hind legs (described previously in males of *Aulocara elliottii*; Bromenshenk & Anderson 1981). We suggest that this element reflects the internal state of the male, i.e. sexual arousal and readiness to mate (prior to mounting attempts). Limited courtship during pre-mounting was previously attributed to both *S. gregaria* (Popov 1958; Strong & Amerasinghe 1977) and *Locusta migratoria* (Oberlin 1973 cited in Strong & Amerasinghe 1977). Oberlin (1973) suggested that this is a result of the high inter-male competition found under the crowded conditions of a locust swarm.

Stridulation (short-previously referred to as “short burst”, “sharp sounds” or “assault-sounds”; and long-previously referred to as “long sounds”, “long burst”, or “whizzing noises”; Loher 1959; Uvarov 1977) and the hind leg (silent) vibration elements (referred to as “cycling of the hind legs”; Strong & Amerasinghe 1977) are known as major characteristics of male sexual pre-copulatory behavior and have been reported to feature during both pre-mounting and mounting (see also Norris 1954; Laub-Drost cited in Uvarov 1977; Otte 1970). While their role is still not fully resolved, in our current observations they were more frequent during mounting (as also mentioned by Loher, 1959). Overall, in addition to its relatively shorter duration, the mounting stage seems to be the more conserved stage in the locust’s reproductive behavior.

Another intriguing behavioral element during pre-mounting is that ofwing-fluttering. This was previously reported for both sexes of the desert locust during sexual interaction (“stationary wings-fluttering" in Loher 1959, Uvarov 1966, 1977; Njagi & Torto 2002). In other acridids wing fluttering was suggested to have a role in mediating release of male volatile substances in relation to mate finding (Uvarov 1966). However, the role of wing fluttering in relation to sexual behavior in the desert locust has not yet been resolved.

### Female sexual behavior

Female desert locusts demonstrated no clear courtship behavior, and were less dominant than males during the sexual interaction (also reported by Norris 1964; Strong & Amerasinghe 1977; Inayatullah et al. 1994). The sexual behavior of the females was also less stereotypic. Upon encountering a male, female behavior comprisedpalp vibration, antennal movement, searching, and self-grooming. A central characteristic of the female’s behavior during both of the pre-copulatory stages was that of hind leg vibration (Loher 1959; Strong & Amerasinghe 1977; Uvarov 1977). During pre-mounting, leg vibration was mostly low amplitude, with less frequent intermitetenthigh amplitude vibration. This is in accordance with Loher’s contention (1959) that the amplitude of this element reflects the level of excitement of the locust (although, the role of this behavioral element in both sexes is still uncertain). It was also suggested that the female’s vibration of her hind legs may serve as a defensive response against the male’s mounting attempts. This is in accord with our major finding, suggesting that the most prominent behavioral elements demonstrated by the females were those related to rejection of the males.

### Sexual conflict

In this study we paid particular attention to the behavior of females and males at the points of possible conflict, preceding mate selection/decision. We suggest two points at which the conflict between the sexes is manifested (traffic lights in Fig 3): the first occurs during pre-mounting and the second during the mounting stage. The first point of conflict may actually appear repeatedly before a male’s attempts to mount the female, and is manifested in two elements: (1) the female’s walking away (“running away” in Loher 1959), and (2) jumping away (Popov 1958; also referred to as “leaping away” in Strong and Amerasinghe,1977). Jumping away better expresses rejection as it frequently followed dislodgment of the male. We did not include kicking during pre-mounting, although intuitively it may serve as a primary rejection element, because kicking is a common reflexive response of locusts, of both sexes, to tactile stimuli by other locusts, regardless of sex (Norris 1962; Siegler & Burrows 1986).

When attempting to mount the female, males displayed two behavioral elements: (1) climbing (described in this work for the first time), or (2) jumping (the more dominant behavior, previously referred to as “attempt to copulate”, “sexual attack”, “copulation attack” or “assault”; Uvarov 1928; Husain & Mathur 1946 cited in Popov 1958; Loher 1959; Pener 1967; Otte 1970). These two elements were often preceded by peering or scanning (lateral swaying of the body from side to side). In both larvae and adult locusts this behavior is related to estimating distance (Kennedy 1945; Wallace 1959). Though not necessarily related to sexual interactions, scanning plays an important role in the premounting stage, serving the males when jumping, and also in the females’ rejection response to an approaching male.

Although, as mentioned above, the display of short stridulation was not very frequent during premounting, its appearance was commonly associated with dislodgement of the male by the mounted female (in agreement with Loher 1959). Based on their differential relative appearance during premounting and mounting, our findings suggest different functional roles for the short and the long stridulation. The overall role of auditory signaling in the courtship behavior of the male desert locust, although previously considered as relatively insignificant (Loher 1959; Keuper et al.1985; Robinson and Hall 2002) would thus appear to be worth revisiting.

In the second point of conflict, during the mounting stage, the interactions between the sexes were more complex. The females used both, direct and indirect rejection elements. Direct rejection comprised jumping and kicking (defensive reaction, Loher 1959), commonly performed immediately after the male had mounted the female, and often presented sequentially, promoting dislodgement of the male from the female’s back (repulsing the male, Loher 1959). In response to the female’s kicking behavior, a few males displayed defensive behavioral elements, including avoidance and blocking. These latter two elements, described here for the first time, may have a major role in assisting the male to overcome female rejection.

The indirect rejection by the female (passive phase, Strong & Amerasinghe 1977), comprising her abdominal bending and abdominal grounding (the latter described here for the first time), is of special interest as it drew a distinctive response from the male: i.e. pressing his hind legs firmly to the ground in an attempt to lift the female.

We examined the efficacy of female jumping and kicking in successfully rejecting males at this conflict point by preventing the females from using their hind legs. Constraining the females indeed resulted in fewer mounting attempts and increased male mounting success. Hence we can safely postulate that a major component of mate-choice by the female is based on consistent and vigorous rejection by way of jumping and kicking. Males, however, overcome female rejection mostly by repeated mounting attempts.

Throughout this study we did not detect any clear signal of female receptivity. High receptivity was best demonstrated passively, whereby passive females did not reject the male (Popov 1958). Twisting of the abdomen, suggested by Ballard et al. (1932; cited in Popov 1958) as a display of receptivity, was never observed in the current study. Another issue that has remained unresolved is that of inter-sexual recognition prior to pre-copulatory behavior. Previous reports have suggested mainly visual, but also chemical, signaling as playing a role in mutual recognition between the sexes in the desert locust (Popov 1958; Uvarov 1977; Pener & Shalom 1987; Obeng-Ofori et al. 1993; Franck & Schmidt 1994; Inayatullah et al. 1994; Ely et al. 2006). Our findings support a major role of visual signals, as we observed that rapid movement by the females (fast walking or jumping) appeared to enhance the males sexual stimulation.

### Concluding remarks

A detailed investigation of the sexual and reproductive behavior is a prerequisite for understanding the evolutionary and ecological dynamics of a species (Kirkendall 1983; Thornhill and Alcock 1983). The comprehensive description presented here of the reciprocal interactions between the sexes in the desert locust thus contributes to our understanding of the biology and behavior of this economically significant pest. The described and presentrd ethogram offers a tool with which to compare behavioral similarities and differences among different orthopteran insects (Paranjape 1985), and specifically among locust species. Here we exclusively described the sexual behavior of the desert locust in the gregarious phase. The knowledge acquired in this study and the tools developed for it will be used for a future comparative investigation of locusts in the gregarious and solitary phases, emphasizing the different features of the sexual conflict in relation to the phase phenomenon.

As noted, the desert locust is one of the most notorious agricultural pests. Major efforts have been invested in investigating the sexual behavior of pest insects (Walgenbach & Burkholder 1987; Rojas et al. 1990; Zahn et al. 2008), with the rationale being that a better understanding of their sexual and reproductive behavior will contribute to the application of pest management (Boake et al. 1996; Suckling 2000). This work may thus also assist in identifying novel targets and generating environmentally friendly methods for locust control.

## Acknowledgments

This work was funded by a grant from the Israel Ministry of Agriculture and Rural Development (891-0277-13)

## Author Contribution

YG performed the experiments, prepared figures and tables. YR prepared figures and tables. AH AA and YG conceived and designed the experiments. All authors wrote and reviewed drafts of the paper

## References

Amerasinghe FP (1978a) Pheromonal effects on sexual maturation, yellowing, and the vibration reaction in immature male desert locusts (Schistocerca gregaria). J Insect Physiol 24(4):309-314. doi: 10.1016/0022-1910(78)90028-8

Amerasinghe FP (1978b) Effects of JHI and JH III on yellowing, sexual activity and pheromone production in allatectomized male Schistocerca gregaria. J Insect Physiol 24(8):603-611. doi: 10.1016/0022-1910(78)90123-3

Ariel G, Ayali A (2015) Locust collective motion and its modeling. PLoS Comput Biol 11(12), p.e1004522. doi: 10.1371/journal.pcbi.1004522

Ayali A, Lange AB (2010) Rhythmic behaviour and pattern-generating circuits in the locust: key concepts and recent updates. J Insect Physiol 56(8):834-843. doi: 10.1016/j.jinsphys.2010.03.015

Ayali A, Yerushalmi Y (2010) Locust research in the age of model organisms: Introduction to the Special issue in honor of MP Pener’s 80th birthday. J Insect Physiol 56(8):831-833. doi: 10.1016/j.jinsphys.2010.05.010

Ayali A, Zilberstein Y, Cohen N (2002) The locust frontal ganglion: a central pattern generator network controlling foregut rhythmic motor patterns. J Exp Biol 205(18):2825-2832.

Baker TC, Cardé RT (1979) Courtship behavior of the oriental fruit moth (Grapholitha molesta): experimental analysis and consideration of the role of sexual selection in the evolution of courtship pheromones in the Lepidoptera. Ann Entomol Soc 72(1):173-188. doi: 10.1093/aesa/80.1.78

Ballard E, Mistikawi AM, Zoheiry MS EL (1932) The Desert Locust, Sehistoeerca gregaria Forsk., in Egypt. Bull. Minist. Agric. Egypt, Cairo, no. 110, 49 pIs.

Berkowitz A, Laurent G (1996) Central generation of grooming motor patterns and interlimb coordination in locusts. J Neurosci 16(24):8079-8091.

Boake CR, Shelly TE, Kaneshiro KY (1996) Sexual selection in relation to pest-management strategies. Annu Rev Entomol 41(1):211-229. doi: 10.1146/annurev.en.41.010196.001235

Bromenshenk JJ, Anderson NL (1981) A study of the mating behavior of Aulocara elliotti (Thomas)(Orthoptera: Acrididae). Trans Am Entomol Soc 107(3):229-247.

Brown MB (1974) Identification of the sources of significance in two-way contingency tables. J Appl Stat:405-413.

Burrows M (1996) The neurobiology of an insect brain: Oxford University Press. New York.

Castrovillo PJ, Cardé RT (1980) Male codling moth (Laspeyresia pomonella) orientation to visual cues in the presence of pheromone and sequences of courtship behaviors. Ann Entomol Soc Am 73(1):100-105. doi: 10.1093/aesa/73.1.100

Charlton RE, Cardé RT (1990) Behavioral interactions in the courtship of Lymantria dispar (Lepidoptera: Lymantriidae). Ann Entomol Soc Am 83(1):89-96. doi: 10.1093/aesa/83.1.89

Chauvin R (1941) Contribution a I’etude physiologique du criquet peterin et du dtterminisme des phenomcnes gregaires. Ann Sot Enr Fr 110:133-272.

Cozzie LR, Irby WS (2010) Anti-insect defensive behaviors in equines post-West Nile virus infection. J Vet Behav 5(1), pp.13-21. doi: 10.1016/j.jveb.2009.08.001

Curkovic T, Brunner JF, Landolt PJ (2006) Courtship behavior in Choristoneura rosaceana and Pandemis pyrusana (Lepidoptera: Tortricidae). Ann Entomol Soc Am 99(3):617-624. doi: 10.1603/0013-8746(2006)99[617:CBIC RA]2.0.CO;2

Eggers A, Gewecke M (1993) The dorsal rim area of the compound eye and polarization vision in the desert locust (Schistocerca gregaria). In: Wiese K, Gribakin FG, Popov AV, Renninger G (ed) Sensory systems of arthropods. Birkhuser, Basel, pp 101’109

EL-Bashir E, Inayatullah C, Ahmed AO, Abdelrahman HE (1993) Models for estimation of population density of late instar nymphs and fledglings of the desert locust, Schistocerca gregaria (Forsk.). Int J Pest Manage 39(4):467’470.

Francke W, Schmidt GH (1994) Potential of semiochemicals for locust control. In: Rembold H, Benson JA., Franzen H, Weickel B, Schulz F (ed) New Strategies for Locust Control. Bonn, Germany: AASTAF, pp.48-52

Gaertner BE, Ruedi EA, McCoy LJ, Moore JM, Wolfner MF, Mackay TF (2015) Heritable variation in courtship patterns in Drosophila melanogaster. G3 (Bethesda) 5(4):531-539.

Haynes KF, Birch MC (1984) The periodicity of pheromone release and male responsiveness in the artichoke plume moth, Platyptilia carduidactyla. Physiol Entomol 9(3):287-295. doi: 10.1111/j.1365-3032.1984.tb00711.x

Hamilton AG (1955) Parthenogenesis in the desert locust (Schistocerca gregaria Forsk.) and its possible effect on the maintenance of the species. Physiol Entomol 30(7-9):103-114. doi: 10.1111/j.1365-3032.1955.tb00188.x

Haskell PT (1957) Stridulation and associated behaviour in certain Orthoptera. 1. Analysis of the stridulation of, and behaviour between, males. The British Anim Behav 5(4):139-48. doi: 10.1016/S0950-5601(57)80020-3

Haskell PT (1958) Stridulation and associated behaviour in certain Orthoptera. 2. Stridulation of females and their behaviour with males. Anim Behav 6(1):27-42. doi: 10.1016/0003-3472(58)90005-8

Husain MA, Mathur CB, Roonwal ML (1946) Studies on Schistocerca gregaria (Forskål) XIII. Food and feeding habits of the desert locust. Indian J. Ent, 8:141-163.

Inayatullah C, El Bashir S, Hassanali A (1994) Sexual behavior and communication in the desert locust, Schistocerca gregaria (Orthoptera: Acrididae): sex pheromone in solitaria. Environ Entomol 23(6):1544-1551. doi: 10.1093/ee/23.6.1544

Injeyan HS, Tobe SS (1981) Phase polymorphism in Schistocerca gregaria: reproductive parameters. J Insect Physiol 27(2):97-102. doi: 10.1016/0022-1910(81)90115-3

Kasuya, E., 1983. Behavioral Ecology of Japanese Paper Wasps, Polistes spp. (Hymenoptera: Vespidae). II. Ethogram and lnternidal Relationship in P. chinensis antenna/is in the Founding Stage. Z. Tierpsychol 63: 303-317. doi: 10.1007/BF02530848

Kennedy JS (1945) Observations on the mass migration of desert locust hoppers. Ecol Entomol 95(5):247-262. doi: 10.1111/j.1365-2311.1945.tb00262.x

Keuper A, Otto C, Latimer W, Schatral A (1985). In: Kalmring K and Elsner N (ed) Acoustic and Vibrational Communication in Insects, Airborne sound and vibration signals of bushcrickets and locusts; their importance for the behaviour in the biotope. Paul Parey. Berlin and Hamburg. pp. 135-142

Kirkendall LR (1983) The evolution of mating systems in bark and ambrosia beetles (Coleoptera: Scolytidae and Platypodidae). Zool J Linnean Soc 77(4):293-352. doi: 10.1111/j.1096-3642.1983.tb00858.x

Klein AL, De Araújo AM (2010) Courtship behavior of Heliconius erato phyllis (Lepidoptera, Nymphalidae) towards virgin and mated females: conflict between attraction and repulsion signals?. J Ethol 28(3):409-420. doi: 10.1007/s10164-010-0209-1

Latchininsky A, Sword G, Sergeev M, Cigliano MM, Lecoq M (2011) Locusts and grasshoppers: behavior, ecology, and biogeography. Psyche 1:1-4 doi: 10.1155/2011/578327

Laub-Drost I (1959) Verhaltensbiologie, besonders ausdrucksausserungen (einschliesslich lautausse-rungen) einiger wanderheuschrecken und anderer orthopteren (Orthoptera: Acrididae: Catantopinae und Oedipodinae). Stuttg Beitr Naturkd 30:1-27

Laub-Drost, I (1960) Verhaltensweisen im Zustand niederer Aktivität bei einigen Wanderheuschrecken und anderen Acridiern (Orthopt.). Ethol 17(5):614-626.

Leonard SH, Ringo JM (1978) Analysis of Male Courtship Patterns and Mating Behavior of Brachymeria intermedia. Ann Entomol Soc Am 71(6):817-826. doi: 10.1093/aesa/71.6.817

Loher W (1959) Contributions to the study of the sexual behaviour of Schistocerca gregaria Forskål (Orthoptera: Acrididae). Proc R Entomol Soc A 34(4-6):49-56). doi: 10.1111/j.1365-3032.1959.tb00237.x

Loher W (1961) The chemical acceleration of the maturation process and its hormonal control in the male of the desert locust. Proceedings of the Royal Society of London B 153(952):380-397. doi: 10.1098/rspb.1961.0008

Mahamat H, Hassanali A, Odongo, H, Torto B, El-Bashir ES (1993) Studies on the maturation-accelerating pheromone of the desert locust Schistocerca gregaria (Orthoptera: Acrididae). Chemoecology, 4(3-4):159-164. doi: 10.1007/BF01256551

Markow TA, Hanson SJ (1981) Multivariate analysis of Drosophila courtship. Proc Natl Acad Sci 78(1):430-434.

Morse AP (1896) Some Notes on Locust Stridulation. J N.Y Entomol Soc 4(1):16-20.

Nishide Y, Tanaka S (2012) Yellowing, morphology and behaviour in sexually mature gynandromorphs of the desert locust Schistocerca gregaria. Physiol Entomo 37(4):379-383. doi: 10.1111/j.1365-3032.2012.00854.x

Njagi PG, Torto B (2002) Evidence for a compound in Comstock-Kellog glands modulating premating behavior in male desert locust, Schistocerca gregaria. J Chem Ecol 28(5):1065-1074. doi: 10.1023/A: 1015222120556

Norris MJ (1954) Sexual Maturation in the Desert Locust (Schistocerca gregaria) Forskal with Special Reference to the Effects of Grouping. Anti-Locust Research Centre.

Norris MJ (1962). Group effects on the activity and behaviour of adult males of the desert locust, Schistocerca gregaria (Forskål) in relation to sexual maturation. Anim. Behav. 10:275-291. doi: 10.1016/0003-3472(62)90051-9

Norris MJ, Richards OW (1964) Accelerating and Inhibiting Effects of Crowding on Sexual Maturation in Two Species of Locusts. Nature, 203:784-785. doi: 10.1038/203784b0

Oberlin UP (1974) Verhaltensbiologische Studien an der europaeischen Wanderheuschrecke Locusta Migratoria L. Entomol Gesell Basel 23:12-23.

Odhiambo TR (1966) Growth and the hormonal control of sexual maturation in the male desert locust, Schistocerca gregaria (Forskål). Trans R Ent Soc Lond 118(13):393-412

Obeng-Ofori D, Torto B, Hassanali A (1993) Evidence for mediation of two releaser pheromones in aggregation behaviour of gregarious desert locust, Schistocerca gregaria (Forskal) (Orthoptera: Acrididae). J Chem Ecol 19:1665-1676. doi: 10.1007/BF00982299

Obeng-Ofori D, Njagi PG., Torto B., Hassanali A, Amiani H (1994) Sex differentiation studies relating to releaser aggregation pheromones of the desert locust, Schistocerca gregaria. Entomol Exp Appl 73(1):85-91. doi: 10.1111/j.1570-7458.1994.tb01842.x

O’shea M (1970) The antennal cleaning reflex in the desert locust, Schistocerca gregaria (Forsk.). Proc. Int. Study Conf. Current and Future Problems of Acridology, London:55’59.

Otte D (1970) A comparative study of communicative behaviour in grasshoppers. Misc. Publ. Mus. Zool. Univ. Mich. 141:1’168.

Ould Ely S, Mahamat H, Njagi, PG, Omer Bashir M, El-Tom El-Amin S, Hassanali A (2006) Mate location mechanism and phase-related mate preferences in solitarius desert locust, Schistocerca gregaria. J Chem Ecol 32(5):1057-1069. doi: 10.1007/s10886-006-9045-8

Paranjape SY (1985) Behavioural analysis of feeding and breeding in Orthopteran insects. Proc Indian Acad Sci 94(3):265-282.

Pener MP (1965) On the influence of corpora allata on maturation and sexual behaviour of Schistocerca gregaria. J. Zool 147(2):119-136.

Pener MP (1967a) Comparative studies on reciprocal interchange of corpora allata between males and females of adult Schistocerca gregaria (Forskdl) (Orthoptera: Acrididae). Proc R Entomol Soc 42(10-12):139-148. doi: 10.1111/j.1365-3032.1967.tb00805.x

Pener MP (1967b) Effects of allatectomy and sectioning of the nerves of the corpora allata on oöcyte growth, male sexual behaviour, and colour change in adults of Schistocerca gregaria. J Insect Physiol 13(5):665-684. doi: 10.1016/0022-1910(67)90117-5

Pener MP, Lazarovici P (1979) Effect of exogenous juvenile hormones on mating behaviour and yellow colour in allatectomized adult male desert locusts. Physiol Entomol 4(3):251-261. doi: 10.1111/j.1365-3032.1979.tb00202.x

Pener, MP, Shalom U (1987) Endocrine manipulations, juvenile hormone and ontogenesis of male sexual behaviour in locusts. Insect Biochem 17(7):1109-1113. doi: 10.1016/0020-1790(87)90130-2

Pener MP, Simpson SJ (2009) Locust phase polyphenism: an update. Adv Insect Physiol 36:1-272. doi: 10.1016/S0065-2806(08)36001-9

Pickford R, Gillott C (1972) Courtship behaviour of the migratory grasshopper Melanoplus sanguinipes (Othoptera: Acrididae). Can Entomol 104(5):715-722.

Popov GB (1958) Ecological studies on oviposition by swarms of the Desert Locust (Schistocerca gregaria Forskal) in eastern Africa. Anti-Locust Bull (31):1-70

Riede K (1987) A comparative study of mating behaviour in some neotropical grasshoppers (Acridoidea). Ethol 76(4):265-296. doi: 10.1111/j.1439-0310.1987.tb00689.x

Robinson DJ, Hall MJ (2002) Sound signalling in Orthoptera. Adv Insect Physiol 29:151-278. doi: 10.1016/S0065-2806(02)29003-7

Roffey J, Popov G (1968) Environmental and behavioural processes in a desert locust outbreak. Nature 219(5153):446-450.

Rojas JC, Malo EA, Gutierrez-Martinez A, Ondarza RN (1990) Mating behavior of Triatoma mazzottii Usinger (Hemiptera: Reduviidae) under laboratory conditions Ann Entomol Soc Am 83(3):598-602. doi: 10.1093/aesa/83.3.598

Rowell CF (1971) Antennal cleaning, arousal and visual interneurone responsiveness in a locust. J Exp Biol 55(3), pp.749-761.

Schmeling F, Wakakuwa M, Tegtmeier J, Kinoshita M, Bockhorst T, Arikawa K, Homberg U (2014) Opsin expression, physiological characterization and identification of photoreceptor cells in the dorsal rim area and main retina of the desert locust, Schistocerca gregaria. J Exp Biol 217(19):3557-3568. doi: 10.1242/jeb.108514

Seidelmann K, Warnstorff K (2001) Ein kombiniertes Y-T-Olfaktometer für Biotests mit grösseren Insekten. Mitt Dtsch Ges Allg Angew Entomol 13:403-408.

Shaluf MI. (2007) An overview on disasters. Disaster Prevention and Management: An International Journal, 16(5):687-703. doi.org/10.1108/09653560710837000

Siegler MV, Burrows M (1986) Receptive fields of motor neurons underlying local tactile reflexes in the locust. J Neurosci 6(2):507-513.

Singh C, Singh J (1977) Rann of Kutch-a trap for the locusts. Indian Journal of Entomology 39(1):23-28.

Suckling DM (2000) Issues affecting the use of pheromones and other semiochemicals in orchards. Crop Prot 19(8):677-683. doi: 10.1016/S0261-2194(00)00090-9

Steedman A (1988) Locust handbook 2nd ed. Overseas Development. Natural Resource Institute London.

Strong L, Amerasinghe FP (1977) Allatectomy and sexual receptivity in females of Schistocerca gregaria. J Insect Physiol 23(1):131-135. doi: 10.1016/0022-1910(77)90118-4

Thornhill R, Alcock J (1983) The evolution of insect mating systems. Cambridge: Harvard University Press.

Uvarov BP (1928) Locusts and Grasshoppers. A Handbook for their Study and Control.

Uvarov BP (1966) Grasshoppers and Locusts: A Handbook of General Acridology. Vol. 1, Anatomy, Physiology, Development, Phase Polymorphism, Introduction to Taxonomy. Published for the Anti-Locust Research Centre at the University Press.

Uvarov BP (1977) Grasshoppers and locusts. A handbook of general acridology Vol. 2. Behaviour, ecology, biogeography, population dynamics. Centre for Overseas Pest Research.

Wajc E, Pener MP (1969) The effect of the corpora allata on the mating behavior of the male migratory locust, Locusta migratoria migratorioides [R. & F.]. Isr J Zool 18(2-3):179-192.

Walgenbach CA, Burkholder WE, Curtis MJ, Khan ZA (1987) Laboratory trapping studies with Sitophilus zeamais (Coleoptera: Curculionidae). J Econ Entomol 80(4):763-767. doi: 10.1093/jee/80.4.763

Wallace GK (1959) Visual scanning in the desert locust Schistocerca gregaria Forskal In J exp Biol.

Wang Q, Millar JG (2000) Mating behavior and evidence for male-produced sex pheromones in Leptoglossus clypealis (Heteroptera: Coreidae). Ann Entomol Soc Am 93(4):972-976. doi: 10.1603/0013-8746(2000)093[0972:MBAEFM]2.0.œ;2

Whitman DW (1990) Grasshopper chemical communication. In: Chapman R and Joem A (ed) Biology of Grasshoppers. New York. John Wiley. pp.357-391

Weis-Fogh T (1956a) The flight of locusts. Sci Am 194:116-126.

Weis-Fogh T (1956b) Biology and physics of locust flight. II. Flight performance of the desert locust (Schistocerca gregaria). Phil. Trans. R. Soc. B: Biological Sciences, 239(667):459-510. doi: 10.1098/rstb. 1956.0008

Wood D, Ringo JM, Johnson LL (1980) Analysis of courtship sequences of the hybrids between Drosophila melanogaster and Drosophila simulans. Behav Genet 10(5):459-466. doi: 10.1007/BF01073650

Wybrandt GB, Andersen SO (2001) Purification and sequence determination of a yellow protein from sexually mature males of the desert locust, Schistocerca gregaria. Insect Biochem Mol Bio 31(12):1183-1189. doi: 10.1016/S0965-1748(01)00064-9

Zahn DK, Girling RD, McElfresh JS, Cardé RT, Millar JG (2008) Biology and reproductive behavior of Murgantia histrionica (Heteroptera: Pentatomidae). Ann Entomol Soc Am 101(1):215-228. doi: 10.1603/0013-8746(2008)101[215:BARBOM]2.0.CO;2

